# Excellent agreement between automated deep learning-based and manual DWI infarct volume measurements in hyperacute stroke

**DOI:** 10.64898/2025.12.17.695031

**Authors:** Yuki Sakamoto, Junya Aoki, Yuji Nishi, Sotaro Shoda, Michika Sakamoto, Takehiro Katano, Junki Yoshimura, Sae Ueda, Takashi Shimoyama, Satoshi Suda

## Abstract

**Background:** Diffusion-weighted imaging (DWI) lesion volume and infarct growth are important imaging markers in acute ischemic stroke, but manual volume measurement is time-consuming and resource-intensive. Deep learning (DL)-based automated segmentation may facilitate rapid assessment; however, its performance on hyperacute DWI has not been sufficiently assessed. The aim was to evaluate the agreement between DL-based automated and manual infarct volume measurements and to compare their ability to predict clinical outcomes in patients treated with mechanical thrombectomy (MT).

**Method:** Consecutive MT-treated patients (September 2014–December 2019) who underwent DWI at admission and at approximately 24 hours were retrospectively analyzed. Manual infarct volume was measured by stroke neurologists. Automated measurements were obtained using DL-based software. Agreement was assessed using Pearson’s correlation, Bland-Altman analysis, and intraclass correlation coefficients (ICC 2,1). Inter– and intra-rater reliabilities were evaluated in a randomly selected subgroup of 150 patients. Predictive ability for a good outcome at 3 months (modified Rankin Scale score 0–2 or stable/improved from premorbid status) was compared using C-statistics and DeLong’s test.

**Results:** A total of 371 patients (677 DWI scans) were included. Manual and automated measurements showed very strong correlation (r = 0.96) with minimal bias (–1.77 mL). The ICC for manual-automated agreement was 0.959 (95% CI, 0.952–0.965), comparable to inter– and intra-rater ICCs. Agreement remained high across onset-to-imaging times and lesion sizes. Predictive abilities for a good outcome were similar for manual and automated admission DWI volume (C-statistics 0.867 vs. 0.861) and infarct growth (0.859 vs. 0.853). Manual follow-up DWI volume showed slightly better predictive ability than automated measurement (0.880 vs. 0.866).

**Conclusion:** DL-based automated infarct volume measurement shows excellent agreement with experienced clinicians, with predictive performance comparable to manual assessment. Automated DWI-based quantification is reliable and feasible for use in hyperacute stroke management.

## Introduction

Hyperintense signals on diffusion-weighted imaging (DWI) reflect cytotoxic edema^1^ and are considered to represent the ischemic core in patients with acute ischemic stroke. DWI hyperintensity shows a physiological change due to ischemic insult, and the extent of DWI hyperintensity is an indicator for reperfusion therapies.^2,3^ In addition, DWI lesion volume (“infarct volume”) or its change, known as infarct growth, is a strong imaging predictor for clinical outcomes,^4^ and, therefore, several recent studies have used infarct growth during the acute phase as a primary outcome.^5,6^ Although infarct volume and infarct growth are important and promising imaging features in acute stroke management, manual volume measurement is highly time-consuming and requires substantial human resources. Obtaining infarct volume or infarct growth in a timely manner in the hyperacute setting is therefore challenging. Software-based approaches can support this task. Some applications estimate infarct volume using an apparent diffusion coefficient (ADC) threshold, but the performance of segmenting DWI hyperintensity solely on a single ADC threshold varies greatly among individuals,^7^ and lesion volume can differ hugely across software even when the same ADC threshold is used.^8^ Moreover, ADC threshold-based software often underestimates or even fails to detect small lesions.^7,9,10^ Recently, fully automated ischemic lesion segmentation algorithms using deep learning (DL) techniques have been developed, and several studies have reported promising results.^9–14^ However, most of these studies used subacute DWI or ADC scans for both training and validation. Although infarct volume or infarct growth can be used as an indication for hyperacute treatment or adjunctive therapy, DL-based software has shown poorer performance for infarct volume on hyperacute DWI than on subacute DWI in prior reports.^11,13^ The aim of the present study was to evaluate the agreement between DL-based automated measurement and manual measurement of infarct volume and infarct growth on hyperacute DWI, in consecutive patients treated with mechanical thrombectomy (MT).

## Methods

From September 2014 through December 2019, consecutive patients with acute stroke treated with emergent MT at admission were retrospectively recruited from a prospective registry.^15,16^ This population was selected because most patients had hyperacute (<6 h) DWI on admission (before MT) and also underwent follow-up MRI approximately 24 h after admission.^16^ Patients who had contraindications to MRI (e.g., cardiac pacemakers), who were treated with MT after admission (e.g., for symptom progression), or who received ≥2 MT procedures during the first 7 days from admission, were excluded. For the purpose of evaluating software performance in real-world settings, DWI scans with any type of artifact (e.g., from metals or motion) were not excluded from this study. MRI, including DWI, was performed on admission and approximately 24 hours later using a commercially available echo planar instrument operating at 1.5 T (Echelon Oval, Hitachi Medical Systems, Tokyo, Japan) in our institution. DWI was carried out using the following parameters: TR/TE, 6000/65 ms; b-values, 0 and 1000 s/mm^2^; field of view, 24 cm; acquisition matrix, 128 × 128; and slice thickness, 4.5 mm, with a 2.5-mm intersection gap. For patients initially scanned at local hospitals, DWI obtained on scanners from the local hospital (magnetic field strength: 0.5 to 1.5 T) was used.

### Manual infarct volume measurement

Infarct volume was measured manually on admission and on follow-up DWI using image analysis software (3D Slicer, http://www.slicer.org). 3D Slicer, built through support from the National Institutes of Health, is a free, open-source, extensible software application for medical image analysis. The patients’ Digital Imaging and Communications in Medicine (DICOM) data were imported, and hyperintense lesions on DWI were semi-automatically outlined on each slice using segmentation tools (Supplemental Figure S1).^15,16^ The hyperintense area was multiplied by the slice thickness plus intersection gap, resulting in a volume measurement (mL). Scans were divided into three sets, and two board-certified stroke neurologists (Y.S. and J.A.) and one neurology resident (Y.N.) independently performed manual measurements. For the present study, 50 patients from each rater’s dataset were randomly selected, and Y.S. (certified neurologist 1) re-measured the infarct volume manually, to assess inter-rater (with J.A. and Y.N.) and intra-rater agreements. The repeated measurements were performed after an interval of approximately 13 months.

### Automatic infarct volume measurement

Patients’ raw DICOM data were loaded into JLK DWI (JLK Inc., Seoul, South Korea), and the infarct volume was analyzed. No pre-processing procedure was applied. We primarily used the automated segmentation results derived from b=1000 images for comparison with manual measurements. The software also provides auxiliary segmentation maps based on the Apparent Diffusion Coefficient (ADC) to support diagnosis (Supplemental Figure S1).

### Statistical analysis

The relationship between manual and automated measurements was examined using a scatterplot and Pearson’s correlation coefficient, and the systematic difference between the two methods was evaluated with Bland-Altman analyses. Because we hypothesized that DWI scans obtained at local hospitals (often low-strength magnetic field) and the presence of symptomatic intracerebral hemorrhagic (sICH) are associated with poor agreement, these cases were highlighted in the analyses. To explore factors associated with discrepancies between manual and automated infarct volume measurements, a multivariable linear regression model was constructed. The dependent variable was the relative infarct volume difference, defined as (|volume_manual_ – volume_automated_|) / {(mean of volume_manual_ and volume_automated_) + 1}. The value 1 was added to the denominator to avoid division by zero. Sex, age, onset-to-imaging time, MRI at local hospital, and sICH were included as independent variables. The intraclass correlation coefficient (ICC) (2, 1) between manual and automated measurements was calculated for all infarct volumes (both on admission and on follow-up). Next, ICCs (2, 1) were evaluated for the randomly selected subgroup of 150 patients (50 patients each from three reviewers) to assess inter– and intra-rater agreements. Manual-automated ICCs (2, 1) were also calculated for this subgroup. ICCs (manual-automated and inter-/intra-rater) were reported by reviewer and scan timing (on admission or 24-h follow-up) as well. Finally, the predictive ability for a good functional outcome at 3 months was evaluated using C-statistics (area under the receiver-operating characteristic curve) for manual and automated infarct volume and infarct growth. A good outcome was defined as a 3-month modified Rankin scale (mRS) score of 0-2 for patients with pre-morbid mRS score ≤2, and as an mRS score that was stable or improved relative to the pre-morbid mRS score for those with pre-morbid mRS score >2. A multivariable binary logistic regression model was constructed with known predictors for good outcome after MT, such as sex, age, hypertension, pre-morbid mRS score, initial National Institutes of Health Stroke Scale (NIHSS) score, successful recanalization (modified Thrombolysis in Cerebral Infarction score ≥2b), onset-to-recanalization time, sICH (SITS-MOST criteria^17^), and initial serum glucose level. The C-statistics were compared pairwise using DeLong’s test.

All statistical analyses were performed using R (Version 4.5.1, R Foundation for Statistical Computing, Vienna, Austria) with the packages *ggplot*, *gtsummary*, *irr*, and *pROC*. Results were considered significant at p<0.05.

## Results

Of 442 acute endovascular therapies performed during the study period, 406 were emergent MTs. Of them, 18 were excluded because they had ≥2 EVTs during the first 7 days from admission, and 11 patients were excluded because they had no MRI examinations due to contraindications. Finally, 377 MTs (225 males [60%], median age 76 [IQR 68-83] years, median NIHSS score 17 [10-23]) for 371 patients were included in the present study (Supplemental Figure S2). The clinical characteristics of the included patients are shown in Table 1. A total of 677 DWI scans (371 on admission and 306 on follow-up) were used to test agreement. MRI on admission was performed at a median 134 (81-350) minutes from stroke onset, and the time from initial to follow-up MRI was 22 (17-26) hours.

**Table 1.** Clinical characteristics of included patients.

| Variables | Total<br>(n = 377) |
| --- | --- |
| Male sex | 225 / 377 (60%) |
| Age (years) | 76 (68-83) |
| Body mass index (kg/m<sup>2</sup>)<sup>a</sup> | 22.3 (19.9-24.5) |
| Hypertension | 237 / 377 (63%) |
| Dyslipidemia | 130 / 377 (34%) |
| Diabetes mellitus | 76 / 377 (20%) |
| Current smoker | 139 / 377 (37%) |
| Atrial fibrillation | 189 / 377 (50%) |
| Premorbid mRS | 0 (0-1) |
| SBP on admission (mmHg) | 158 (135-177) |
| DBP on admission (mmHg) | 87 (77-100) |
| NIHSS score on admission | 17 (10-23) |
| Onset-to-first MRI (min)<sup>b</sup> | 134 (81-350) |
| Site of arterial occlusion |  |
| ICA | 88 / 377 (23%) |
| MCA M1 | 144 / 377 (38%) |
| MCA M2 or distal | 102 / 377 (27%) |
| ACA | 4 / 377 (1.1%) |
| VA | 6 / 377 (1.6%) |
| BA | 25 / 377 (6.6%) |
| PCA | 8 / 377 (2.1%) |
| rt-PA treatment | 130 / 377 (34%) |
| Effective reperfusion | 314 / 377 (83%) |
| Onset-to-Reperfusion time (min) | 262 (190-470) |
| Symptomatic ICH | 29 / 377 (7.7%) |
| Stroke etiology |  |
| CE | 216 / 377 (57%) |
| LAA | 75 / 377 (20%) |
| Others | 86 / 377 (23%) |
| Glucose (mg/dL)<sup>c</sup> | 126 (110-154) |
| Manual DWI volume on admission (mL)<sup>b</sup> | 11 (3-41) |
| Manual DWI volume on follow-up (mL)<sup>d</sup> | 20 (6-67) |
| Time from initial to follow-up MRI, (hr)<sup>e</sup> | 22 (17-26) |

Data are shown as number/total number (%) or median (interquartile range).
mRS indicates modified Rankin Scale; SBP, systolic blood pressure; DBP, diastolic blood pressure; NIHSS, National Institute of Health Stroke Scale; MRI, magnetic resonance imaging; ICA, internal carotid artery; MCA, middle cerebral artery; ACA, anterior cerebral artery; VA, vertebral artery; BA, basilar artery; PCA, posterior cerebral artery; rt-PA, recombinant tissue-plasminogen activator; ICH, intracerebral hemorrhage; CE, cardioembolism; LAA, large-artery atherosclerosis; DWI, diffusion-weighted imaging.
a Missing in 9 cases.
b Missing in 5 cases due to no MRI on admission
c Missing in 2 cases
d Missing in 72 cases due to no follow-up MRI
e Missing in 77 cases due to no MRI either on admission or follow-up

The correlation between manual and automated infarct volume measurements was very strong (Pearson’s correlation coefficient r = 0.96, p < 0.01, Figure 1A), even in cases with small infarct volumes (enlarged panel in Figure 1A). Bland-Altman analysis showed that the mean difference (bias) was –1.77 mL, with 95% limits of agreement of –33.92 to 30.38 mL (Figure 1B). These very strong correlations and small mean differences were consistent across various times from stroke onset (or last known well time for unwitnessed patients) to MRI imaging (Supplemental Figure S3). On multivariable linear regression analysis, male sex (standardized coefficient –0.16, 95% CI –0.32 to –0.01), MRI performed at a local hospital (0.18, 0.10 to 0.25), and the presence of sICH (0.12, 0.02 to 0.23) were independently associated with larger relative differences between manual and automated measurement (Supplemental Table S1). Visually, slightly larger mean differences were observed in cases with MRI scans at a local hospital within 3 hours from stroke onset on Bland-Altman analysis (Supplemental Figure S3A).

**Figure 1.**
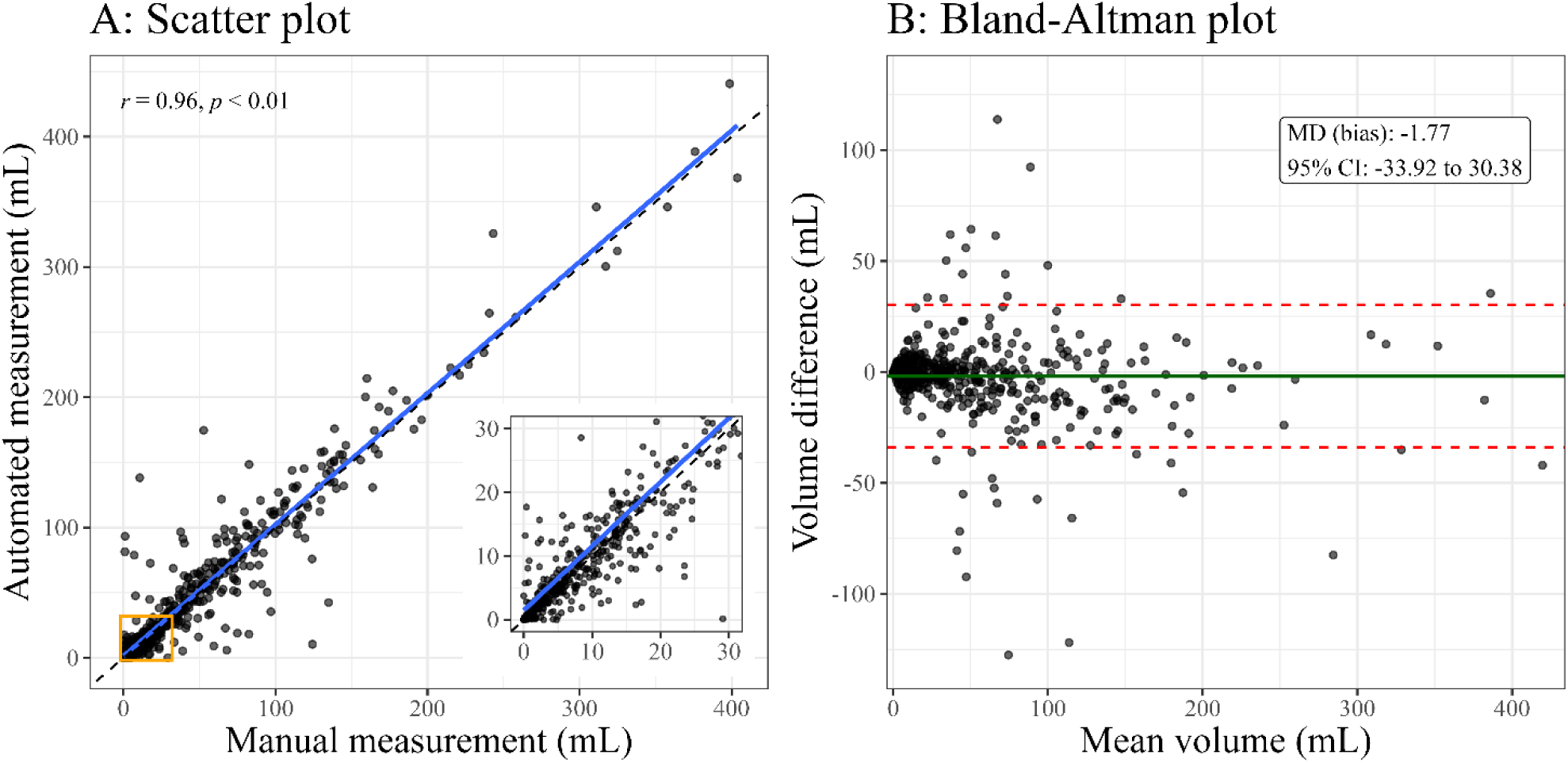
Correlation of and systematic difference between manual and automated infarct volume measurements. (A) Scatterplot indicates a very strong (Pearson’s r = 0.96) correlation. The solid blue line represents the linear best fit line, and the dashed black line represents the line of identity (y = x). The lower-right panel enlarges the region with infarct volume <30 mL (enclosed in an orange square in the main plot). (B) Bland-Altman plot. The green line indicates the mean volume difference, and the dashed red lines indicate its 95% confidence interval (limits of agreement). MD indicates mean difference; CI, confidence interval

**Figure 2.**
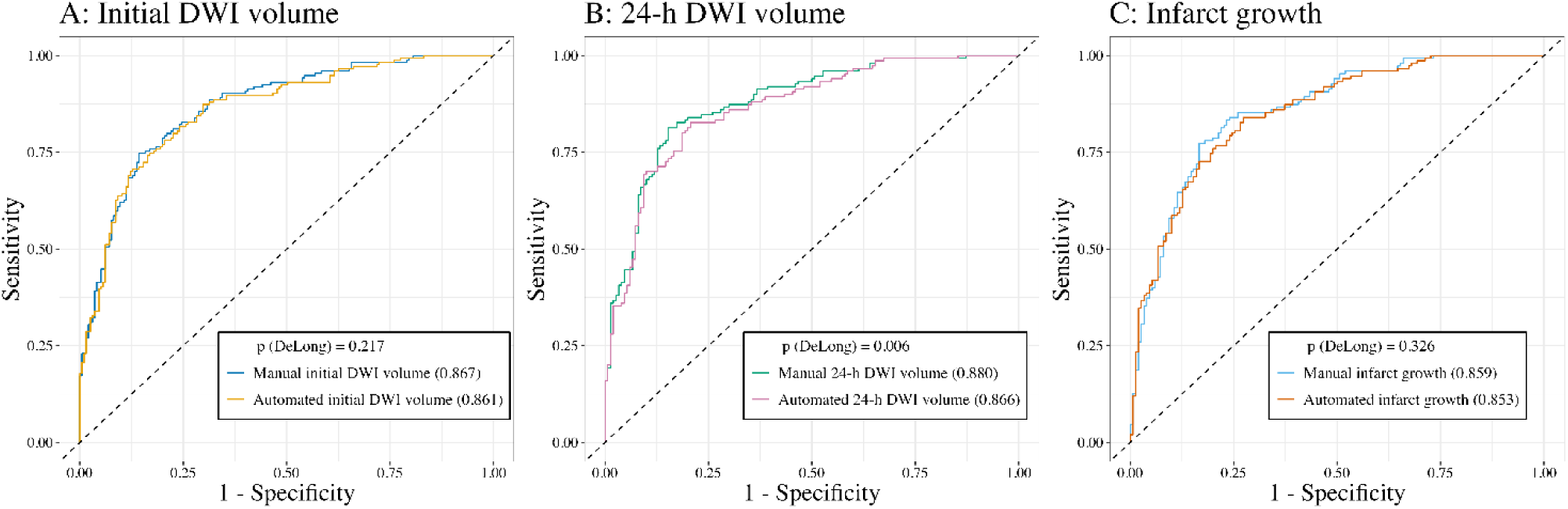
Predictive ability for 3-month good outcome using manual and automated volume measurements. Receiver operating characteristics curves are shown for multivariable binary logistic regression models including each volume measurements, in addition to known predictors for good outcome after MT, such as sex, age, hypertension, pre-morbid mRS score, initial NIHSS score, successful recanalization, onset-to-recanalization time, sICH, and initial serum glucose level. Numbers in parentheses within the legend represent C-statistics. DWI indicates diffusion-weighted imaging; MT, mechanical thrombectomy; mRS, modified Rankin Scale; NIHSS, National Institutes of Health Stroke Scale; sICH, symptomatic intracerebral hemorrhage.

ICC (2, 1) between manual and automated infarct volume measurements was 0.959 (95% confidence interval [CI] 0.952 to 0.965) for all DWI scans (Table 2). The ICC values were numerically higher for follow-up DWI scans (0.971, 0.964 to 0.977) than for the initial scan (0.935, 0.920 to 0.947).

**Table 2.**
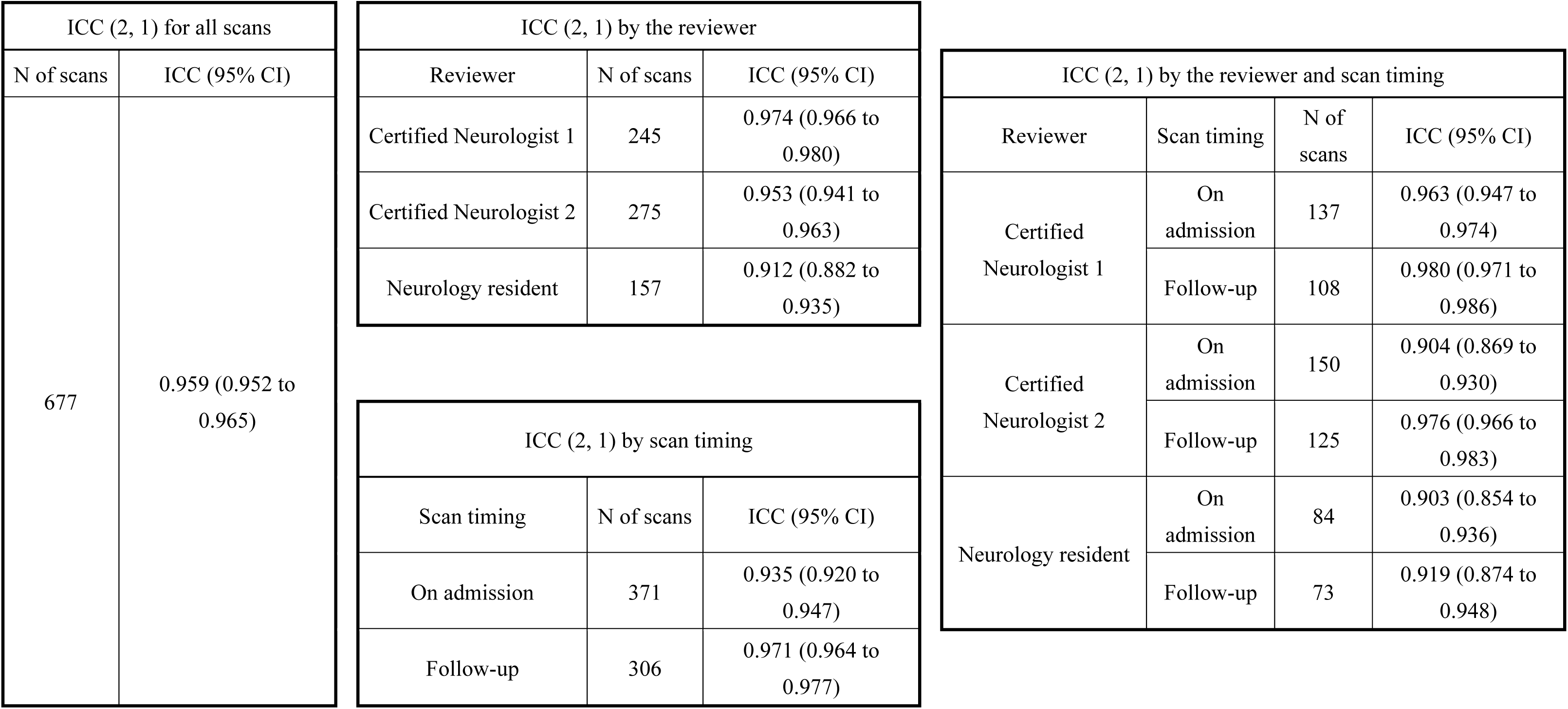
Intraclass correlation coefficient between manual and automated infarct volumes for all scans.

The 150 randomly selected patients with 275 DWI scans were similar to the non-selected patients, except for a higher prevalence of dyslipidemia (Supplemental Table S2). The agreements between manual and automated measurements were also excellent in this subgroup. The ICC (2, 1) values for manual-automated agreement were almost the same as those for inter-rater and intra-rater agreements among experienced neurologists (Table 3).

**Table 3.** Comparison of ICCs between automated and manual measurements and human inter-/intra-rater reliabilities.

| N of scans | ICC (95% CI) |  |
| --- | --- | --- |
|  | Neurologist 1 vs. Automated | Neurologist 1 vs. Neurologist 1 |
| 92 | 0.953 (0.930 to 0.969) | 0.945 (0.917 to 0.963) |
|  | Neurologist 2 vs. Automated | Neurologist 2 vs. Neurologist 1 |
| 92 | 0.948 (0.923 to 0.965) | 0.963 (0.945 to 0.976) |
|  | Resident vs. Automated | Resident vs. Neurologist 1 |
| 92 | 0.928 (0.893 to 0.952) | 0.921 (0.883 to 0.947) |

| Scan timing | N of scans | ICC (95% CI) |  |
| --- | --- | --- | --- |
|  |  | Neurologist 1 vs. Automated | Neurologist 1 vs. Neurologist 1 |
| On admission | 50 | 0.963 (0.935 to 0.979) | 0.943 (0.903 to 0.967) |
| Follow-up | 42 | 0.947 (0.903 to 0.971) | 0.944 (0.898 to 0.969) |
|  |  | Neurologist 2 vs. Automated | Neurologist 2 vs. Neurologist 1 |
| On admission | 49 | 0.934 (0.886 to 0.962) | 0.972 (0.952 to 0.984) |
| Follow-up | 43 | 0.957 (0.922 to 0.977) | 0.954 (0.917 to 0.975) |
|  |  | Resident vs. Automated | Resident vs. Neurologist 1 |
| On admission | 50 | 0.962 (0.934 to 0.978) | 0.944 (0.903 to 0.968) |
| Follow-up | 42 | 0.901 (0.823 to 0.945) | 0.901 (0.823 to 0.946) |

The predictive abilities for good functional outcome at 3 months were similar between manual and automated measurements for initial DWI volume (C-statistics 0.867 vs. 0.861, p = 0.217) and infarct growth (0.859 vs. 0.853, p = 0.326). For follow-up DWI volume (at approximately 24 hours), lesion volume by manual measurement showed better predictive ability than by automated measurement (0.880 vs. 0.866, p = 0.006, Figure 3).

## Discussion

The present study showed that fully automated infarct volume measurement showed excellent agreement with manual measurement. This high agreement was consistent even for hyperacute DWI scans obtained within the first few hours after stroke onset. The agreement between manual and automated measurement was comparable to both inter-rater and intra-rater agreements among experienced stroke neurologists and a neurology resident. The ability of automated infarct volume or infarct growth to predict good functional outcome at 3 months was similar to that of manual measurement for initial DWI volume and infarct growth.

The excellent agreement between manual and automated DWI lesion volumes is in line with previous reports that showed that a DL-based fully automated segmentation algorithm can segment DWI hyperintensity in high accordance with manual measurement.^9–14^ A novel aspect of the present study is that agreement was validated using consecutive hyperacute DWI scans, obtained as early as a median of 134 minutes from stroke onset. Most of the previous studies trained and validated models using subacute DWI scans. The present results indicate that automated segmentation can perform well even on hyperacute DWI, where only subtle signal changes are present in many cases. The strong agreement and minimum systematic difference (mean difference or bias in the Bland-Altman analysis) were consistent across a wide range of scan timings and lesion sizes. Moreover, the ICC (2, 1) values for manual-automated agreement were almost the same as those for human inter-and intra-rater agreements. These findings indicate that automated segmentation of DWI hyperintensity is reliable, robust, and practical for patients with acute stroke undergoing reperfusion therapy.

The predictive ability for a good outcome at three months using automated infarct volume or infarct growth was similar to that obtained using manual measurement. Given the inherent limitation in reviewing DWI scans that “true” (histopathologically proven, for example) infarct volume at the time of scanning cannot be obtained, an automated segmentation algorithm should be evaluated not only by agreement with human raters, but also by its ability to predict clinically meaningful outcomes. In this context, the present results of slightly but significantly lower predictive performance of automated measurement for follow-up (at 24 hours) DWI volume seems to be interesting; the manual-automated volume agreement is consistently high, but the predictive performance differs. This and the other result that sICH is the independent factor for larger relative volume difference may suggest that human raters can segment better or more meaningfully than an algorithm in complex images, because follow-up DWI scans in MT-treated patients frequently show heterogeneous lesions with hemorrhagic transformation.

The strengths of the present study were including consecutive patients with acute stroke who were treated with MT, available MRI scans in hyperacute phase and follow-up scans at approximately 24 hours, and evaluating long-term clinical outcomes. Moreover, the software was used in a setting different from its development environment, in a different healthcare system in a different country, MRI equipment, and clinical workflow. However, several limitations should be addressed. First, this was not a prospective study. The manual infarct volume measurements were originally performed for other research purposes,^15,16^ and scans from 150 patients were re-evaluated for inter– and intra-rater reproducibilities for the present study. Second, although MRI was performed for almost all patients (99% of the consecutive MT-treated patients without contraindications underwent MRI in this registry^15^), not all patients had initial MRI and 17.5% of the included patients lacked follow-up MRI. Last, because the software has been evaluated only in East Asian cohorts, its generalizability to other populations remains unknown.

In conclusion, automated DWI-based infarct volume measurement using a DL-based algorithm showed excellent agreement with and minimum bias against manual measurement by experienced stroke neurologists. Agreement between the software and human raters was comparable to that among human raters themselves. Automated infarct volume and infarct growth showed predictive performance for long-term functional outcome similar to that of manual measurement.

## Acknowledgements

The authors would like to express their deepest gratitude to all members of the SU and radiology and emergency departments. This work was partly supported by JSPS KAKENHI Grant Number JP25K10778 and Takeda Science Foundation Grant Number 2025043074.

